# Serotype has a significant impact on the virulence of 7th pandemic *Vibrio cholerae* O1

**DOI:** 10.1101/2020.10.16.342279

**Authors:** Stefan L Nordqvist, Kaisa Thorell, Frida Nilsson, Madeleine Löfstrand, Arvid Hagelberg, Jan Holmgren, Michael R. Lebens

## Abstract

Of over 200 different identified *Vibrio cholerae* serogroups only the O1 serogroup is consistently associated with endemic and epidemic cholera disease. The O1 serogroup has two serologically distinguishable variants, the Ogawa and Inaba serotypes, which differ only by a methyl group present on the terminal sugar of the Ogawa O-antigen but absent from Inaba strains. This methylation is catalyzed by a methyltransferase encoded by the *wbeT* gene, which in Inaba strains is disrupted by mutation. It is currently thought that there is little difference between the two serotypes. However, here we show, using isogenic pairs of O1 El Tor *V. cholerae*, that Inaba strains show significantly different patterns of gene expression and are significantly less able than the corresponding Ogawa strains to cause cholera in an infant mouse infection model. Our results suggest that changes in gene expression resulting from the loss of the *wbeT* gene lead to reduced virulence and possibly also reduced survival fitness outside the human host.

**Author Summary:** The bacterium *Vibrio cholerae* causes the pandemic diarrheal disease cholera. Despite many identified serotypes of *V. cholerae* only one, O1, causes pandemic cholera. The O1 serotype of pandemic *V. cholerae* has two distinguishable variants (called Ogawa and Inaba) long considered to be clinically and epidemiologically equivalent. Cholera outbreaks consist only of one the two variants at any time. In general, Ogawa strains cause the majority of outbreaks with relatively short-lived Inaba outbreaks occurring sporadically. We have suggested earlier that Inaba outbreaks occur during periods of environmental selective pressure against the Ogawa serotype. We demonstrate here that the two variants are not clinically equivalent. The Ogawa serotype is better able to respond to infection in an animal model by up regulating the expression of virulence genes essential for disease development. We suggest that this phenomenon is the result of wider ranging differences in gene expression resulting from the mutation that converts Ogawa into Inaba strains, and may help to explain the dominance of the Ogawa serotype in nature.

## Introduction

Cholera is the most severe of all the infectious diarrheal diseases and a major health problem particularly in Asia and sub-Saharan Africa where endemic disease and epidemic outbreaks cause significant morbidity and mortality. It is accepted that cholera is under-reported but even so, there are an estimated 2.86 million cases per year in endemic areas alone resulting in 100,000 deaths, the majority in sub-Saharan Africa(1, 2). The causative agent of cholera is the bacterium *Vibrio cholerae* which produces a powerful enterotoxin (cholera toxin) and infection is spread mainly via contaminated water or food. *V. cholerae* thrives in brackish and estuarine waters worldwide but of over 200 different identified serogroups, only the O1 serogroup is associated with pandemic cholera. Although the O139 serogroup emerged briefly in the 1990s causing widespread outbreaks in Asia, the O1 serogroup caused all seven cholera pandemics that have occurred since the beginning of the 19th century and is effectively the sole cause of cholera worldwide in modern times(2). The ongoing 7^th^ pandemic which started in the 1960s is caused by organisms of the El Tor biotype that share a clonal origin from a common ancestor which appears to have emerged in the early 20th century. The previous six pandemics are thought to have been caused by the classical biotype which is now extinct as a cause of pandemic cholera but which was also of the O1 serogroup(3).

In both *V. cholerae* biotypes, the O1 serogroup has two serologically distinguishable variants, the Ogawa and Inaba serotypes. The only apparent difference between them is the methylation of the terminal sugar of the poly-perosamine O-antigen in Ogawa strains catalyzed by an S-adenosylmethionine (SAM)-dependent methyltransferase encoded by the *wbeT* gene. In Inaba strains, this gene is inactivated by mutation(4, 5). Prevailing opinion suggests that there is no difference between the two serotypes with respect to their ability to cause cholera or to survive in the environment(6, 7). However, worldwide cholera caused by the Ogawa serotype is predominant and recent work suggests that serotype switches may result from selective pressure based on the structure of the O-antigen(8). Whole-genome sequencing of large numbers of clinical isolates shows that Inaba lineages generally tend to emerge and then die out. New Inaba outbreaks when occurring, are caused by Inaba strains arising from mutations in circulating Ogawa strains, suggesting that Ogawa strains are overall fitter than Inaba strains (9) although other than the epidemiological evidence, there is little experimental data to confirm this. We wished to address whether we could observe any differences between the two serotypes that might account for a difference in virulence and/or other fitness manifestation and thereby help to explain the predominance of Ogawa in nature and the persistence of the *wbeT* gene in the O1 serotype. Although indistinguishable in terms of growth in liquid culture, using isogenic variants of Ogawa and Inaba we could demonstrate that there are significant differences in gene expression between the serotypes. Furthermore, Inaba strains are significantly inferior to Ogawa strains in their ability to colonize the infant mouse intestine and to cause diarrhea reflecting significant differences in the expression of key virulence genes. Our results show for the first time that there are underlying differences in gene expression and virulence between the Ogawa and Inaba serotypes that may also go some way to explain the predominance of the Ogawa serotype in nature.

## Results

Two pairs of isogenic strains of El Tor O1 *Vibrio cholerae* were constructed, each pair comprising one Inaba and one Ogawa strain. The first set was derived from strain Phil6973 isolated from a patient in India in 1973 and is currently a component of all prequalified killed oral cholera vaccines. Phil6973 has the Inaba serotype due to a stop codon at position 252 in the *wbeT* gene. It was converted to the Ogawa serotype by replacing the mutant *wbeT* gene with a wild-type gene from the Ogawa strain VX44945 resulting in the strain MS1571. Subsequently, the *wbeT* gene in MS1571 was deleted in order to produce the Inaba strain MS1712, in which the *wbeT* gene was entirely absent. A second pair of isogenic strains was generated from the clinical Ogawa isolate A493 isolated in Bangladesh in 2012. The *wbeT* gene was deleted in a similar fashion to generate the Inaba strain MS1843. However, when the genome of MS1843 was sequenced it was found that apart from the deletion of the *wbeT* gene there was a non-synonymous mutation in the *crp* gene (I52S). The parental strain A493 did not carry this mutation and a new strain MS1972 was constructed in which the *crp* sequence corresponded with the wild type. Subsequent experiments were performed with both *wbeT* deletion strains, MS1843 and MS1972 as well as the original Inaba strain Phil6973.

### Growth comparison in rich medium

When isogenic strains MS1571 and MS1712 were grown LB medium, high salt medium and AKI medium there was no detectable difference in growth between the Ogawa and Inaba variants ((figures 1a, 1b and 1c respectively). Similar results were obtained with the isogenic pairs A493 and MS1972 (Supplementary figure S1). Competition experiments performed *in vitro* also demonstrated that when grown in a co-culture neither variant out-grew the other (data not shown). Growth under AKI conditions induces the expression of virulence genes in 7^th^ pandemic strains of O1 *V. cholerae* (10). We therefore tested whether there were any differences in the expression of CT between corresponding Ogawa and Inaba strains. It was consistently found that Inaba strains expressed higher levels of CT than their Ogawa counterparts. This is shown for strains MS1571 (Ogawa) and MS1712 (Inaba) in figure 1d.

**Fig 1.**
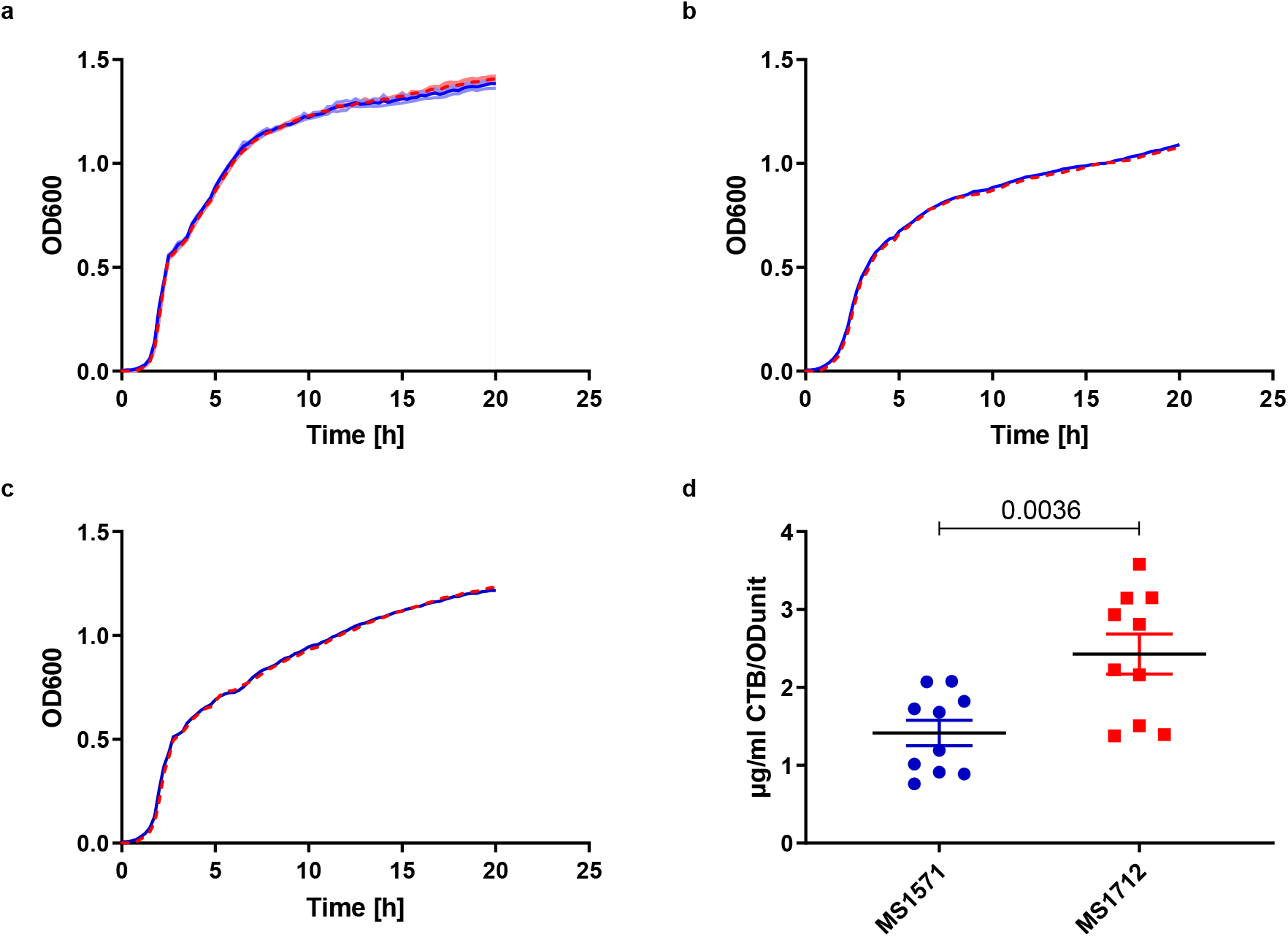
Direct comparison of growth and CTB production of Ogawa and Inaba isogenic strains. The two isogenic strains MS1571 and MS1712 were grown under different conditions A) LB at 37°C, B) LB high salt 30°C, C) AKI medium 30°C. No differences in growth were detected. Ogawa strains in blue and Inaba strains in dotted red. D) In separate experiments levels of CTB production were measured after growth under AKI conditions O/N. Mean +-SEM Two-tailed Unpaired t-test p=0.0036, t=3.345, df=18.

### Transcriptome analysis of isogenic strains

For transcriptomic analysis of the isogenic strains, MS1571 and MS1712 cultures were grown in triplicate in a high salt medium at low temperature. RNAseq analysis showed that under these conditions there were significant differences (P<0.01) in the expression of no less than 472 out of 3519 genes (13,4%) with either higher or lower expression in the Inaba strain when compared with the Ogawa strain (supplementary table S1). Of these, 333 were expressed at higher levels in the Ogawa strain. Nearly all the genes in the purine pathway were expressed at reduced levels in the Inaba strain suggesting a lower requirement for adenosine (figure 2A). Similarly, although genes for methionine synthesis were unaffected, expression of SAM synthetase was expressed at a significantly lower level in the Inaba strain reflecting a lower requirement for SAM in these cells. Among genes previously associated with virulence *CRP*, *toxR*, and *ompT* were all expressed at lower levels in Inaba compared to Ogawa cells. Genes involved in quorum sensing differed between Ogawa and Inaba with small but significant differences in *cqsS* and *aphA* (figures 2B and 3A).

**Fig 2.**
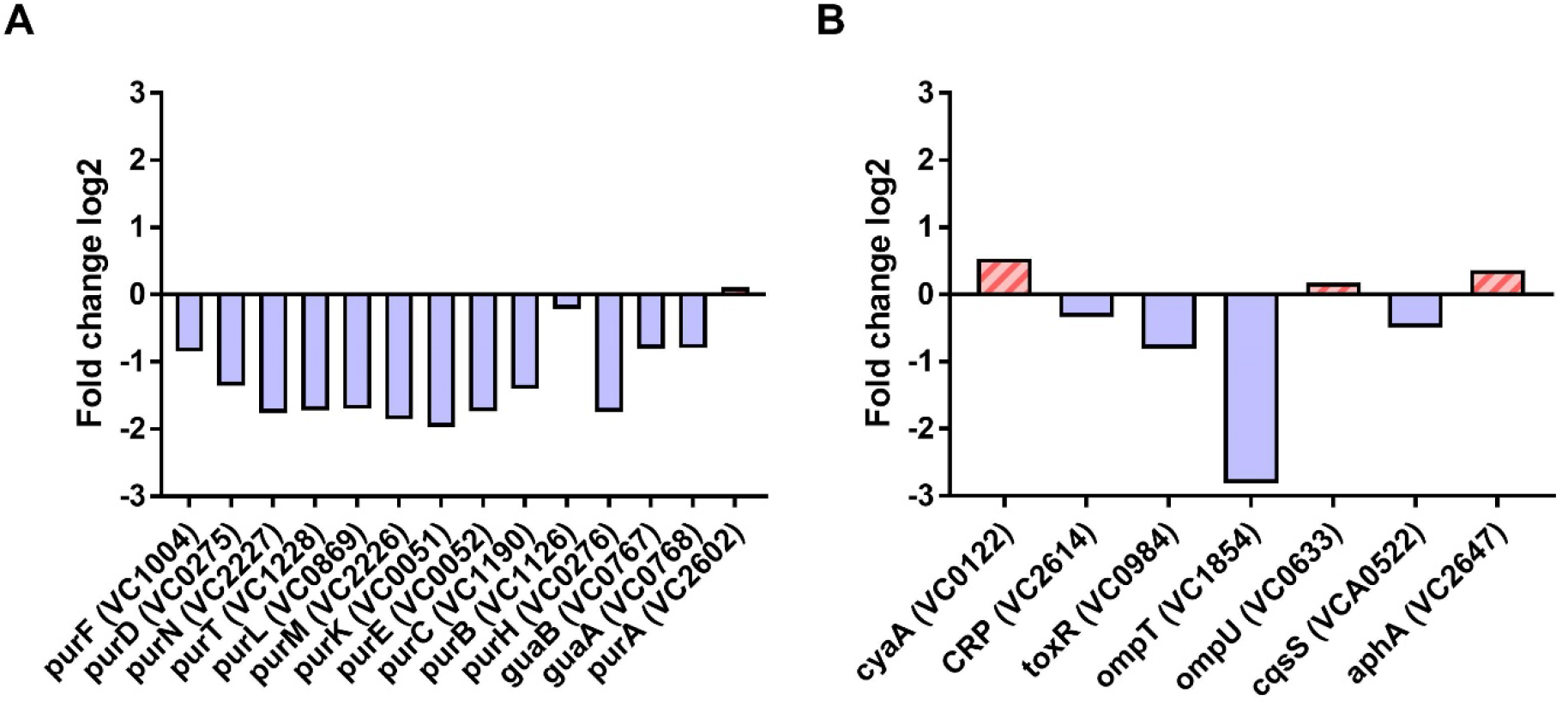
Transcriptomic analysis of the isogenic strains of MS1571 and MS1712. (A) significant differences in expression of genes from the purine synthesis pathway with lower expression in Inaba strains compared to Ogawa strains and (B) observed differences in genes associated with virulence or quorum sensing. Blue bars (negative fold change) represent higher expression in Ogawa and Red striped bars (positive fold change) represent higher expression in Inaba.

**Fig 3.**
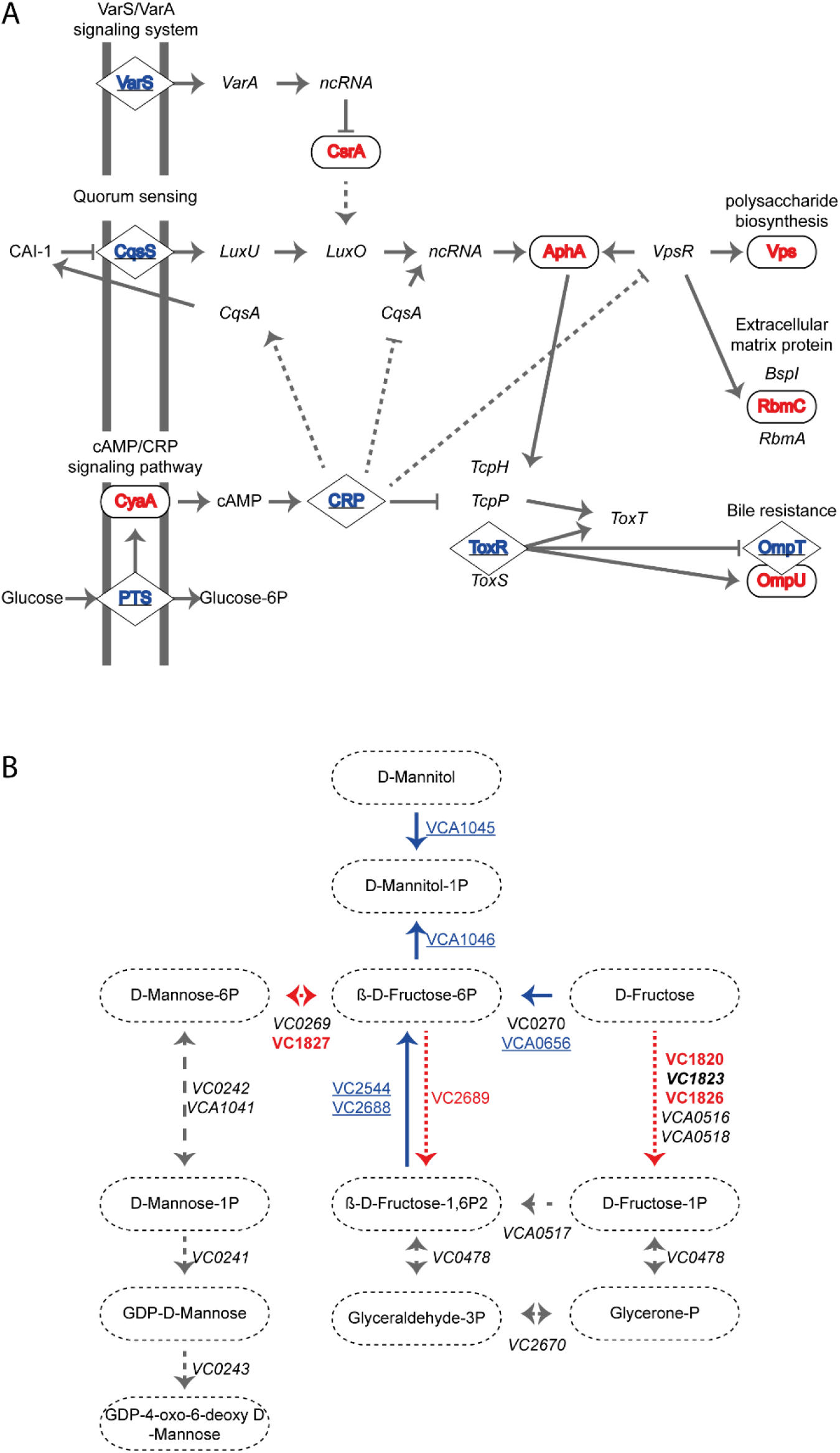
Pathways in which expression of genes differs significantly between Ogawa and Inaba variants of otherwise isogenic strains. RNAseq shows significant changes in expression of genes involved a) in quorum sensing(11), biofilm formation and bile resistance, and b) on fructose and mannose metabolism(12) using the KEGG(13-15) pathways as illustration. Genes with higher expression in Ogawa cells and the reactions their products catalyze are shown in blue and underlined. Genes with higher expression in Inaba cells and the reactions their products catalyze are shown in red. Gene products in bold are also represented in figure 4. Genes with unchanged expression are shown in grey and italic.

Interestingly, a gene that was significantly upregulated in the Inaba strain was *manA* (VC1827), which encodes mannose-6-phosphate isomerase involved in the metabolism of fructose and mannose. On closer inspection, we found that the expression of a number of other genes associated with fructose metabolism were also altered (figure 3B). In addition to *manA*, the associated permease (VC1826) and the regulatory gene controlling their expression (VC1825) were also upregulated in the Inaba strain whilst the surrounding genes also involved in fructose transport were down-regulated (figure 4). Other examples of gene expression reflecting shifts in metabolism are shown in supplementary figures S2 and S3.

**Fig 4.**
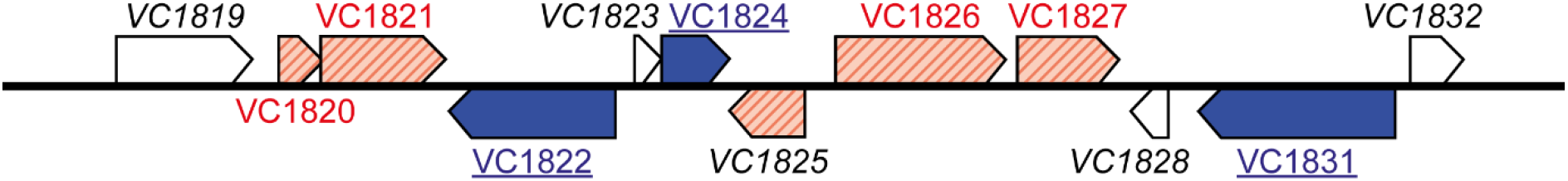
A gene cluster in *Vibrio cholerae* O1 chromosome 1 with genes significantly differentially expressed in Ogawa and Inaba variants of otherwise isogenic strains. The genes affected are involved in fructose metabolism and transport. The roles of VC1820, VC1826, and VC1827 are shown in figure 3B. VC1825 is an *araC* type regulatory gene controlling the expression of VC1826 and VC1827. VC1821 and VC1822 are both involved in fructose uptake. VC1824 and VC1831 are both kinases involved in transmembrane transport. Genes higher expressed in Ogawa (blue), Inaba (red), and genes with no significant difference in white.

### Infection of Infant mice

The results of the transcriptomics showed that there are profound changes in gene expression resulting from deletion of the *wbeT* gene. Some of the involved genes were associated with virulence and might, therefore, impact on the ability of *wbeT* mutants to respond to changes resulting from the introduction into a host intestine. In order to investigate this, the ability of isogenic Ogawa and Inaba strains to multiply and cause cholera in an infant mouse infection model was examined. Groups of mice were infected with the two pairs of isogenic strains and a number of criteria were used to assess the severity of infection and disease. The bacteria were administered together with a blue dye, and the first striking visible difference between the groups was the amount of stool-associated stain in the cages after 20 hours of observation, figure 5. The Ogawa strains clearly gave more staining than the same dose of the corresponding Inaba bacteria indicating more diarrhea. A ten-fold increase in the infecting dose of Inaba bacteria resulted in increased diarrheal staining which still did not reach the level obtained with the low-dose Ogawa infection at the same time-point.

**Fig 5.**
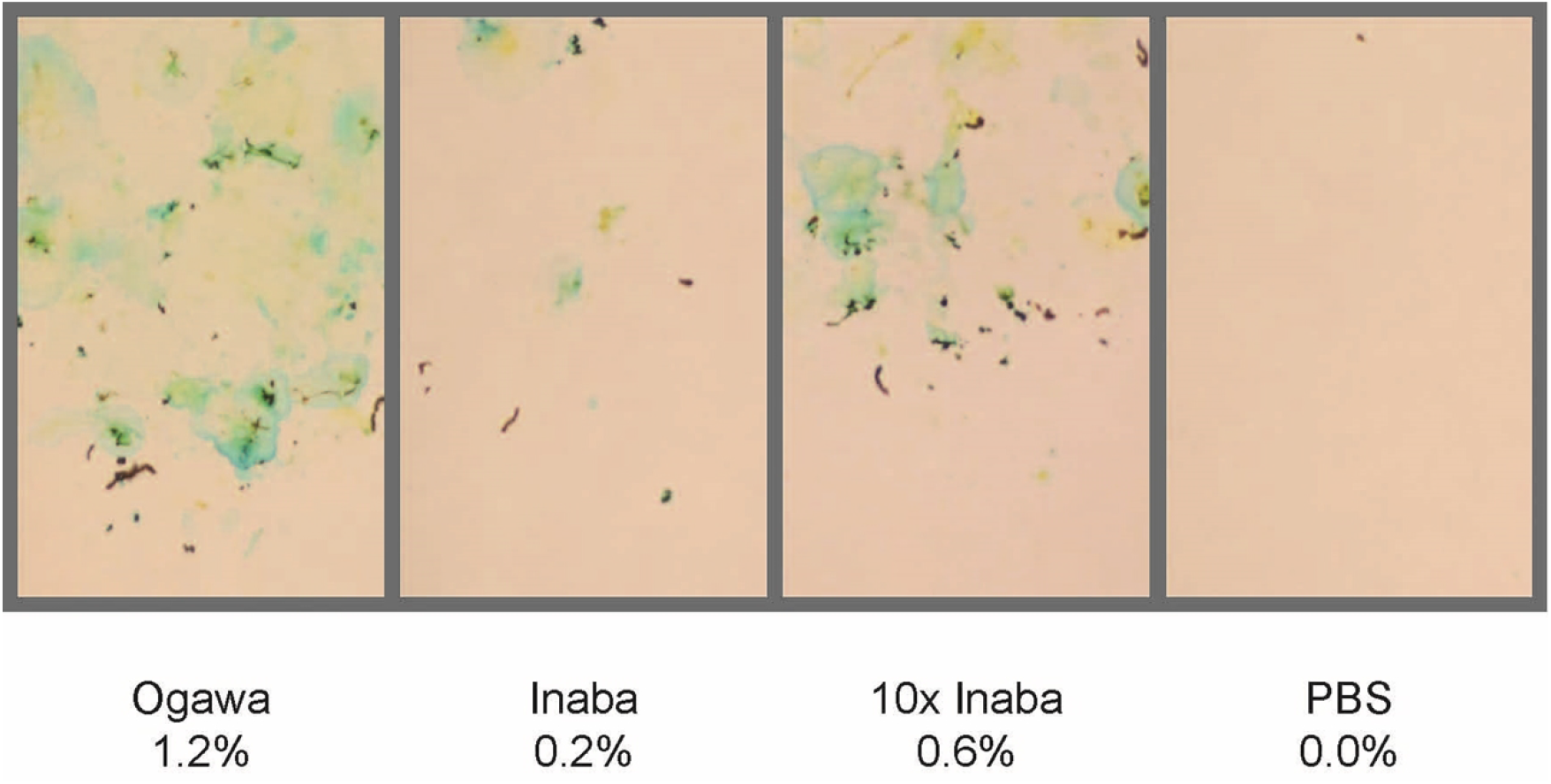
Bedding showing staining due to diarrhea from infected infant mice. Representative images of staining on bedding paper from groups of 12 infant mice infected with either Ogawa, Inaba, 10x dose Inaba, or PBS. Percentages refer to the area covered by the blue stain on the bedding paper determined as described in the materials and methods.

The significant difference between the same infecting dose of Ogawa and Inaba bacteria is also evident for isogenic strains when using more established criteria for cholera infection and disease in this model; weight loss, the weight ratio of the intestine to the carcass, and the number of *Vibrio cholerae* bacteria in the intestine, (figure 6). For the same infection dose, the Ogawa strains (MS1571 and A493) gave more weight loss than the respective Inaba strains (MS1712 and MS1843/MS1972), higher intestine to carcass weight, and more *V. cholerae* in the intestine upon sacrifice. There was no difference between the results obtained in these experiments between the strains MS1843 and MS1972 as shown in supplementary figure S4. Importantly, we found that the Phil6973 strain parental Inaba strain and the *ΔwbeT* strain derived from it were both attenuated when compared to the Ogawa derivative MS1571.

**Fig 6.**
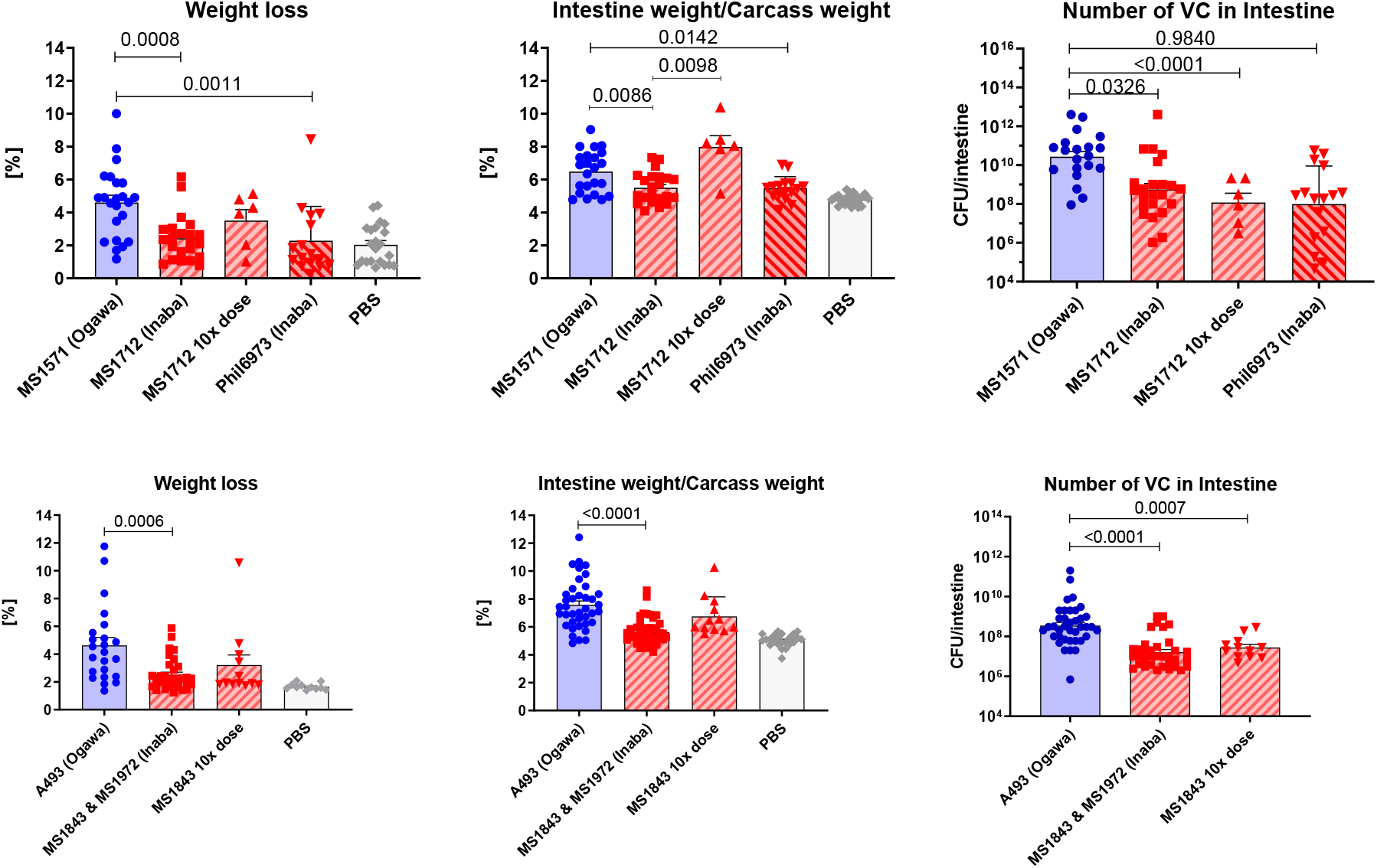
Comparison of disease severity using different criteria in infant mice infected with isogenic strains that were either Ogawa or Inaba. Weight loss, intestine to carcass weight, and the number of *Vibrio cholerae* in the intestine for both sets of isogenic strains. Pooled data from seven different experiments showing mean + SEM and p adjusted values less than 0.05. Total number of mice per group from 7 independent experiments: MS1571 (23), MS1712 (23), MS1712 10x dose (6), phil6973 (16), PBS (23), and A493 (41), MS1843 (32), MS1972(10), MS1843 x10 dose (12), PBS (23). Brown-Forsythe test: F (DFn, DFd) values for: MS1571 vs MS1712 Weight loss 0,7199 (5, 89), Intestine/Carcass 6,438 (5, 90), Number of VC 1,58 (4, 66). A493 vs MS1843 Weight loss 4,424 (3, 75), Intestine/Carcass F 7,485 3, 112), Number of VC F 1,199 (2, 82).

### Virulence gene expression in AKI medium and in infant mice

Having demonstrated that Inaba strains are consistently unable to colonize the infant mouse small intestine as efficiently as the corresponding Ogawa strains it was next tested whether this was dependent upon differences in expression in key virulence genes including *tcpA* and *ctxAB*. This was done by analysis of gene expression of these genes in the intestine of the infected mice using RT-PCR. The results are shown in figure 7. It can be seen that both *tcpA* and *ctxAB* are highly up-regulated in the Ogawa strains in the mouse intestine.

**Fig 7.**
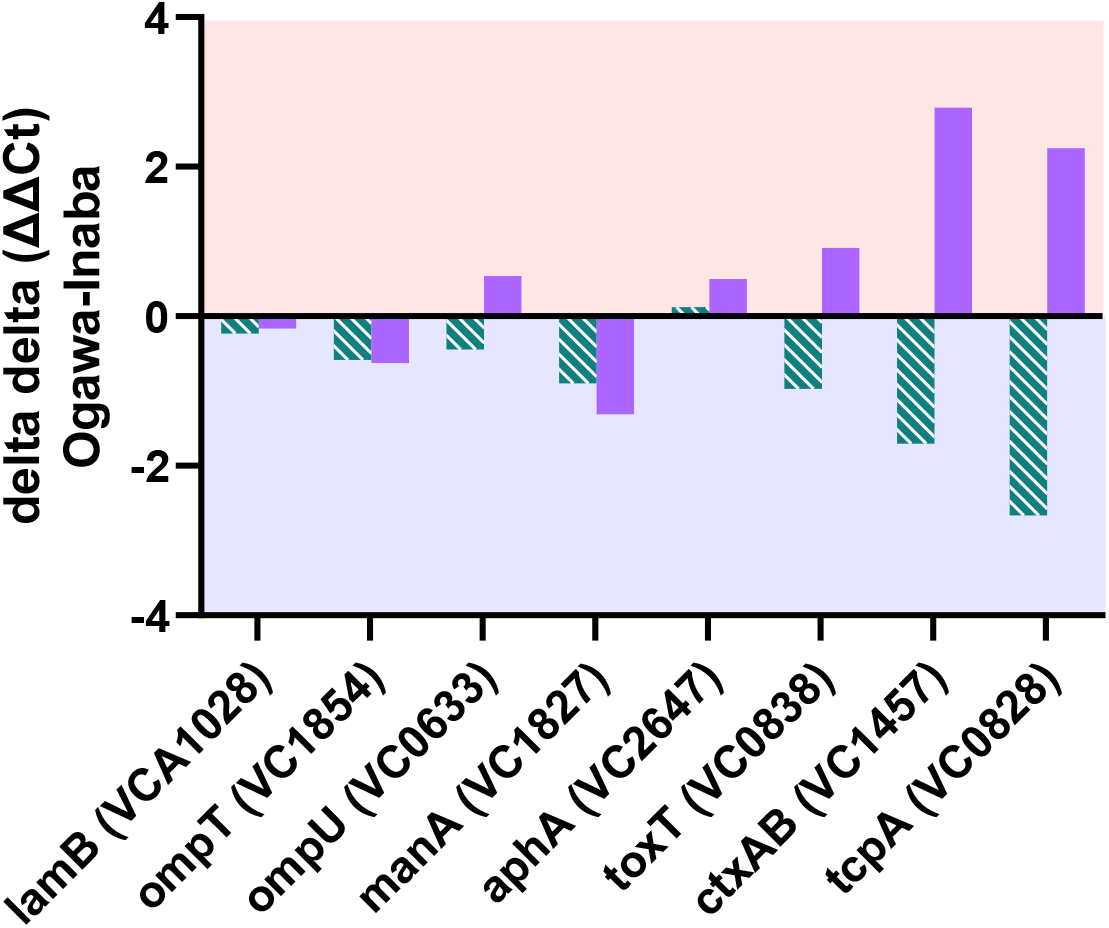
RT-PCR analysis of the isogenic strains of A493 and MS1843 *in vivo* and *in vitro*. Observed differences in expression of genes associated with virulence. ΔΔCT (Ogawa-Inaba) values are shown. Striped green bars and purple bars represent *in vivo* and *in vitro* expression respectively. Negative ΔΔCT values represent higher expression in Ogawa and positive ΔΔCT values represent higher expression in Inaba. *In vivo* groups had five animals per group and *in vitro* had six replicates per group. The results reflect the relative amount of gene expression comparing Inaba with Ogawa strains after normalization against expression of the *recA* gene.

Conversely, the same genes are relatively up regulated in Inaba strains when grown in AKI medium. This is in agreement with differences in expression of CT detected in Inaba strains under full AKI conditions, which as described above, were slightly but consistently higher than those seen in corresponding Ogawa strains.

## Discussion

The observation of two serotypes, Ogawa and Inaba, in clinical isolates of *V. cholerae* O1 is recurrent in outbreaks all over the world but the wider biological implication in terms of colonization and fitness of this switching is not known. Epidemiological evidence suggests that whereas both serotypes cause cholera, the Ogawa serotype persistently spreads causing far more cases than Inaba. Whereas it can be postulated that Ogawa has a better overall fitness than Inaba, no work has been done to determine at what level this is functioning or the cause of periodic serotype switching in endemic areas as exemplified by Karlsson et al. (8) and Baddam et al. (16). The overall similarity of Inaba strains compared to their isogenic Ogawa counterparts in terms of growth and reported virulence have made systematic study of differences between the two serotypes difficult and this is essentially the first study to address this question. Thus, isogenic pairs were generated from clinical isolates of *V. cholerae* O1 7^th^ pandemic strains that were either Ogawa or Inaba through deletion of the *wbeT* gene. Despite no discernable differences when cultures were grown side-by-side in rich medium, there were significant differences in gene expression between the Inaba and Ogawa strains. In all, over four hundred different genes were affected indicating that deletion of *wbeT* has wider implications than merely abrogating methylation of LPS.

The mechanisms underlying all these transcriptional changes are not fully understood but a significant down-regulation of genes involved in purine synthesis and synthesis of SAM in Inaba strains can be attributed to a decreased demand for these molecules compared to Ogawa strains. Moreover, inactivation of a highly active SAM-dependent methyltransferase might be expected to have far-reaching consequences due to changes in ATP and SAM turnover.

Many of the differences in gene expression between Ogawa and Inaba strains could be compensatory adjustments due to altered energy distribution in the cell and, consistent with this, decreases in expression of one gene resulting from the *wbeT* mutation could be seen to be balanced by an increase in expression of another gene with a related function (Figure 5). Further examples are shown in figures S2 and S3. Thus, unchanged growth in rich medium may reflect adjustments in the expression of metabolic pathways that occur to maintain intracellular homeostasis.

Other changes were seen in genes associated with quorum sensing and virulence. These may affect responses to environmental stress that are not apparent under the growth conditions used (17–19) and could be linked to changes in intracellular levels of SAM. SAM is a precursor to a number of important signaling molecules involved in quorum sensing (20, 21) and therefore in the ability of cells to sense and respond to changes in their environment.

The small but significant differences observed in expression of quorum sensing and virulence genes prompted an analysis of CT expression under AKI conditions under which Inaba strains consistently produced slightly higher levels than Ogawa strains. These were the first observed phenotypic differences between the two serotypes and even suggested that Inaba strains might be more virulent than Ogawa strains. The infant mouse model is a well-established cholera infection model that is widely used and mimics acute human disease in several important features including dependence on the toxin co-regulated pilus (TCP) and expression of CT(22). The transition from growth in rich medium to growth in the infant mouse intestine is a dramatic change in environment that imposes significant stress on the bacteria. Inaba cells were much less able to respond to the change than their Ogawa counterparts growing significantly less in the intestine and giving rise to significantly less diarrhea despite producing more CT *in vitro*. This effect was seen in both Inaba strains Phil6973 and the *wbeT* deletion derivative MS1712.

Strikingly, when the expression of *tcpA, ctxAB,* and *toxT* was analyzed in RNA extracted from the intestine of infected mice, it was seen that in all the mice tested, the levels of all three genes were up-regulated, but significantly more so in the mice infected with Ogawa bacteria, with *tcpA* being the most highly up-regulated (figure 7). The higher levels of expression of *ctxAB* by Ogawa cells suggest that despite the *in vitro* results, Ogawa cells produce more CT in *vivo* than Inaba cells. This again suggests that the Inaba bacteria are less able to respond to the changes in environment as they enter the mouse intestine by up-regulating key virulence genes. The results also demonstrate that the AKI growth conditions do not reflect changes in virulence gene expression observed *in vivo*.

The observed differences are unlikely to be entirely due to the difference in the surface LPS but rather reflect other changes in gene expression that result from the loss of the *wbeT* gene. Indeed, in terms of virulence, the differences observed between isogenic Ogawa and Inaba strains are very similar to those seen in *crp* mutants that are significantly attenuated in the infant mouse model (23). CRP is a stress response protein and is one of the genes whose expression was significantly lower in Inaba strains grown under normal laboratory conditions. It was therefore unfortunate that when the *wbeT* gene was initially removed from strain A493, a non-synonymous mutation led to an amino acid change in the *crp* gene. Despite similar results in the Phil6973 derivatives that did not carry this mutation the *wbeT* deletion in A493 was repeated and shown not to carry the mutation in *crp*. The results with this strain were similar and suggest that the *crp* mutation did not play a role in the reduced ability of the strain to infect infant mice (figure S4).

Clearly, much more work needs to be done in order to analyze the gene expression of the different serotypes in the context of the infection model. However, from the results so far we can suggest that in addition to the role of the *wbeT* gene in determining serotype, the impact it has on the distribution of energy and availability of quorum sensing molecules in the cell is likely to impact virulence, affecting how cells respond to stresses encountered in the environment and associated with infection. The loss of *wbeT* in our experiments clearly compromises the severity of disease caused by *V. cholerae* O1. It remains to be seen whether these differences are reflected in the amount of diarrhea and the number of bacteria shed from human individuals infected by the two serotypes. The results are consistent however, with the observed overall superior ability of Ogawa cells to persist and spread compared to Inaba strains.

Since even slight changes to the O1 serotype result in significant changes in the way the organisms interact with their environment, our findings may go some way to explaining the association of the O1 serogroup with cholera over seven pandemics caused by two biotypes in the last 200 years and the predominance of the Ogawa serotype. The ability of O1 *V. cholerae* to cause pandemic cholera may be dependent not only on the O1 serotype but also and possibly primarily on the maintenance of the *wbeT* gene that is responsible not only for the methylated Ogawa serotype but also, as indicated by this study, for promotion of bacterial virulence and perhaps overall survival fitness in nature.

## Material and Methods

### Bacterial strains and plasmids

The bacterial strains used in the current study are shown in Table 1. All strains were maintained on Luria Bertani (LB) plates supplemented when necessary with appropriate antibiotics and stored in a 17% glycerol stock solution at −80°C. Unless indicated otherwise, liquid cultures were grown in LB broth at 37°C as previously described(24).

**Table 1.**
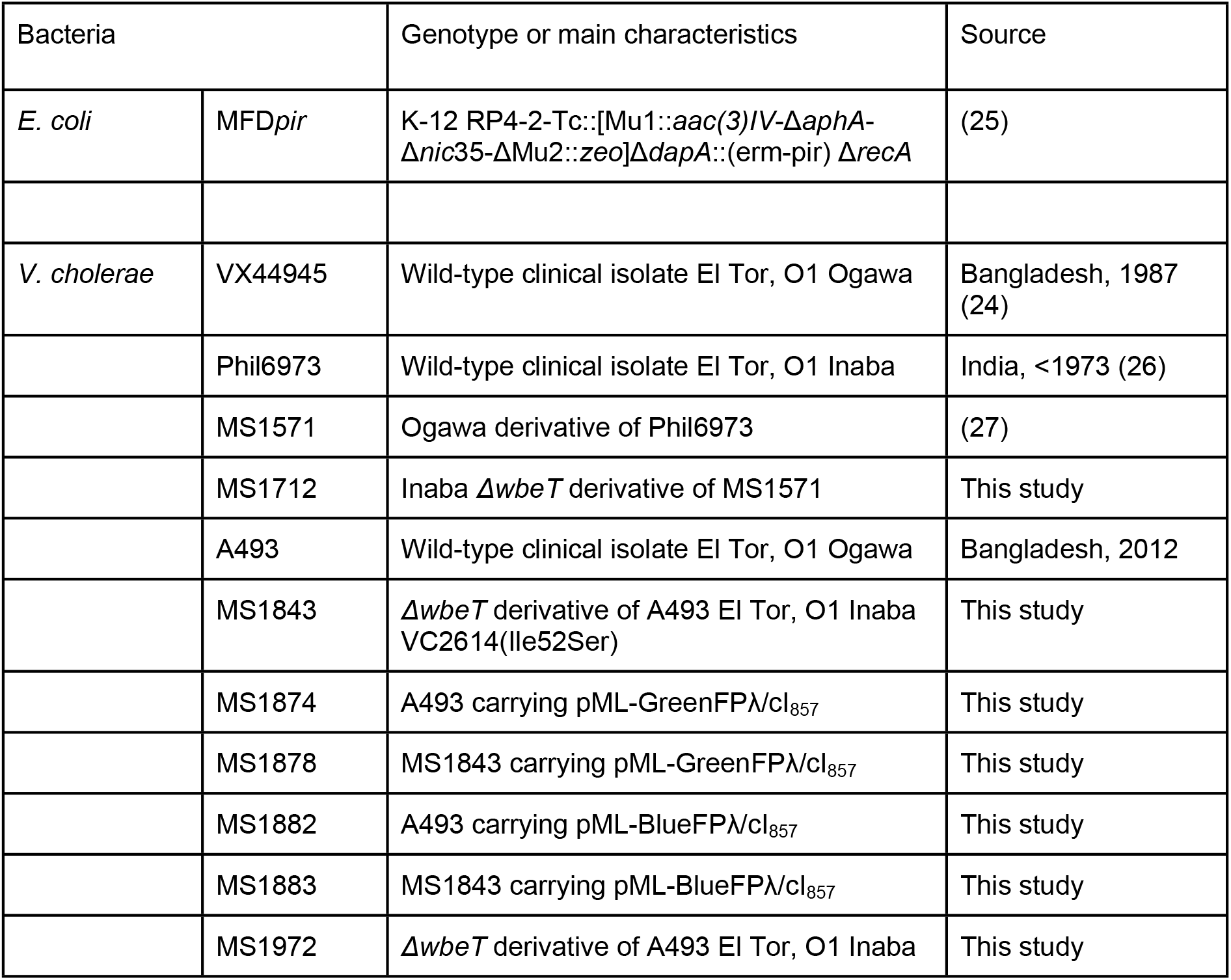
Bacterial strains used in the current study.

The pML-GreenFPλ/cI_857_ and pML-BlueFPλ/cI_857_ were kindly provided by pML-Biokonsult AB (Sweden).

### Constructing Isogenic strains

In order to construct isogenic strains that were Ogawa or Inaba gene replacement was achieved using the suicide vector pMT-suicide/*sacB*(27) (Genebank accession: KF188719.1).

Two fragments flanking the *wbeT* gene were amplified by PCR using primer pairs *wbeT5* (5‘-GGCTTTAGTGAATCGCGATTTGTCGG-3’) with *wbeT_deletion_linker_rev* (5‘-GTCGACGCGGCCGCGATATCACAGAATCAACTTGCAGATGCAGGTTTG-3’) and *wbeVf* (5‘-GGCGTATTACGGTACTACAAGGGTCTAG-3’) with *wbeT_deletion_linker_fwd*(5‘-GATATCGCGGCCGCGTCGACTGCAAGTTCAACAGACATTTCCGAAGAG-3’). The two resulting fragments were combined using primerless PCR, finally amplifying with the primer pair *wbeT5/wbeVf* to generate a fragment in which the *wbeT* gene was deleted. This fragment was inserted into the suicide pMT-suicide/*sacB* and a Km^R^ gene flanked by *lox* sites were inserted between SalI and EcoV sites, figure S5. The final plasmid was used to transform *E. coli* strain MFD*pir* and the resulting strain was used to introduce the plasmid into the recipient strains Phil6973 and A493 conferring resistance to both kanamycin and chloramphenicol. The two strains were then passaged for two days in LB broth supplemented with kanamycin before plating out onto LB agar plates containing no salt but supplemented with sucrose (6% w/v). Individual colonies were picked onto duplicate LB agar plates supplemented with kanamycin and chloramphenicol respectively in order to screen for loss of the plasmid. The resulting strains were Kanamycin resistant and chloramphenicol sensitive. Deletion of the *wbeT* gene was confirmed by sequencing DNA fragments obtained by PCR amplification using the primers *wbeT5* and *wbeVf*. In both generated strains the Km^R^ gene was removed by Cre-mediated recombination. The *cre* gene was introduced using an expression plasmid conferring chloramphenicol resistance in which the Cre expression was induced by addition of IPTG to the growth medium. Cells were grown overnight in LB broth supplemented with chloramphenicol and IPTG to a final concentration of 1 mM. The cells were then serially diluted and spread on LB agar plates to obtain single colonies. These were then picked onto duplicate plates to check for sensitivity to kanamycin. A kanamycin sensitive colony was taken and streaked out onto LB agar. The Cre plasmid was lost due to its inherent instability in the absence of selection with chloramphenicol. The phenotype of the resulting strains was checked by agglutination with an O1 and Ogawa-specific monoclonal antibodies (Fitzgerald, United States)

In the case of Phil6973 the original strain is Inaba and in order to obtain an Ogawa derivative, we used the same suicide plasmid-based procedure to introduce a wild-type *wbeT* gene back into the strain in which the gene had been deleted. In this case, the selection was based solely on the acquisition and loss of chloramphenicol resistance combined with a change in phenotype from Inaba to Ogawa.

### Growth curves

Bacteria were revived from −80°C glycerol stock on LB-Agar plates at 37°C for 16h. 3 colonies were used for inoculation in 5 ml LB medium for 4h at 37°C at 180 rpm. Strains were set to the same OD by diluting with PBS and inoculated either in LB broth, LB broth high salt(28) or AKI(10) medium. The high salt medium was used to maximize growth owing to the halophilic nature of *V. cholerae*. The inoculated media were distributed in a 24 or 48 well cell culture plates (Nunc) and were incubated in a Synergy™ 2 (Biotek, United States) plate reader at 30°C or 37°C, measuring optical density every 15 min at λ=600nm.

### Competitive growth

The growth of isogenic strains in competition experiments was done using strains carrying plasmids pML-GreenFPλ/cI_857_ and pML-BlueFPλ/cI_857_ that express the green fluorescent protein and blue fluorescent protein respectively when induced by incubation at 42°C. The plasmids are essentially identical except for small differences in the structural genes of the fluorescent proteins. Furthermore, the fluorescent proteins are not expressed during the competition experiments since they were performed at 30°C.

Briefly, isogenic pairs of strains, one carrying the pML-GreenFPλ/cI_857_ plasmid and pML-BlueFPλ/cI_857_ were grown up overnight in 5 ml LB broth supplemented with ampicillin (100 μg/ml). The cells were washed and resuspended in PBS and the OD_600_ was adjusted to 0.5. The cells were then mixed in at a ratio of 1:1. Serial dilutions of the suspension were then spread onto LB agar plates supplemented with ampicillin (100 μg/ml) and incubated at 30°C overnight in order to determine the actual number of colony-forming units (CFUs). In order to determine the ratio of Inaba to Ogawa cells in the suspension after growth overnight at 30°C, the plates were transferred to 42°C in order to express the green and blue fluorescent proteins. The actual ratio of one serotype to the other was determined by counting the number of colonies expressing each of the fluorescent proteins.

50 μl of the mixed cell suspension was used to inoculate 5 ml LB broth supplemented with ampicillin and the resulting culture was incubated at 30°C for 14h with shaking (180 rpm). Serial dilutions of the culture were again spread onto LB agar plates in order to determine the total number of CFUs and the ratio of Inaba to Ogawa cells was determined by transferring the plates to 42°C and counting colonies with different fluorescence.

The cultures were then passaged every 14 hours over a period of five days and the ratio of Inaba to Ogawa cells determined as described.

### RNA isolations for RNAseq

Cultures of each of the isogenic strains MS1571 and MS1712 were grown in triplicates in 25 ml high salt LB medium at 23°C in 250 ml Erlenmeyer flasks to OD_600_ = 1.00. RNA extracted using Qiagen RNeasy kit for gram-negative bacteria and cDNA library preparation was performed using the NuGen Ovation stranded RNAseq kit with custom rRNA depletion specific for *V. cholerae* and libraries were prepared from each culture. The sequencing was done using the Illumina HiSeq2000 instrument generating on average 12.6 million reads per sample. The draft genome of *V. cholerae* strain Phil6973 was ordered based on the Vibrio cholerae O1 biovar El Tor N16961 complete genome using Mauve order contig tool(29). The ordered contigs were annotated using the prokka pipeline(30) using a manually curated annotation of the N16961 genome, replacing the split open reading frames VC0255 and VC0256 with a wild-type WbeT protein sequence (GenBank: JF284685.1) as the primary reference source. The annotated genome was used to create a library for the STAR aligner(31) and used as a reference for downstream alignment of the RNA-seq reads. After quality trimming and filtering of the sequence data using TrimGalore(32), the reads were mapped to the Phil6973 reference, as described above and counting of the transcripts was performed using HTSeq-count(33). Statistical analysis and differential expression analysis was performed using the DESeq2 R package(34).

### RNA isolations for RT-PCR

Cultures of each of the isogenic strains A493 and MS1843 were grown on LB plates at 30°C overnight. 5 colonies of each culture were used to inoculate 10ml AKI medium, incubated at 30°C standing still for 4 hours. 2 ml of the culture was centrifuged for 2 minutes at 13’000xg. The pellet was resuspended in 0.5 ml RNA Later (Qiagen, Germany), stored at 4°C overnight and then transferred to −80°C until time for RNA extraction. For the *in vivo* samples; at the time of sacrifice, the intestine was placed in an Eppendorf tube containing 0.5ml RNALater (Qiagen, Germany), cut into small pieces with scissors and immediately placed on ice. The intestine was stored at 4°C O/N and then transferred to −80°C until time for RNA extraction. RNA was extracted using the SV Total RNA Isolation System (Z3100, Promega, USA) for gram-negative bacteria and cDNA library preparation was performed using the GoTaq^®^ 2-Step RT-qPCR System (A6010, Promega, USA). The RT-PCR was done using the Applied Biosystems™ 7500 Real-Time PCR System and primers shown in the supplementary data table S2 (35-38). CT values were normalized using the expression of the housekeeping gene *recA* (VC0543) before comparison of the expression of virulence and quorum sensing genes in the Ogawa and Inaba serotypes.

### Infant mouse infection model

Pregnant female Swiss outbred CD1 mice were purchased from Charles River Laboratories (Germany) and housed at Lab of Experimental Biomedicine (Gothenburg, Sweden). Four days after birth infant mice weighing 3.6±0.9 grams were separated from their mothers, individually marked, and placed at 26°C for 4 hours before being randomly grouped and infected with virulent *V. cholerae* bacteria. Group sized was determined depending on the number of pups born, normally ten mice per group, and never fewer than four.

To maximize the efficiency of infection bacteria were cultured in AKI medium without shaking to an OD_600_ value of approximately 0.5. The cells were then centrifuged 13’300xg for 2 minutes and re-suspended in fresh PBS and the OD_600_ was adjusted to 0.5 whereafter the suspension was diluted 1:20. Blue food dye (E133, Dr. Oetker, Germany) was added to the suspension to a final concentration of 0.05%. The infant mice were orally infected at 1300-1330 by gastric gavage using a non-cutting, round-tip stainless steel feeding needle (AgnTho’s AB, Sweden, 7900 = 25 mm 24G) with 50μl blue suspension (approx. 7.5×10^5^ CFU) or PBS and were kept in the dark in groups at 26°C for 20 hours and sacrificed between 0900-0930 the day after.

All stages of the experiments were blinded in that the staff doing the infections did not know which strain they were infecting with and did not know which group were treated with what when assessing the clinical criteria. Animals that during the course of the experiments showed clinical signs of severe disease i.e. change of skin coloration from pink to blue or grey, labored breathing and persistent recumbency, were immediately euthanized.

The weight of each mouse was measured before the experiment start(w_s_), immediately after infection(w_0_), and 20h after infection(w_20_). Weight loss at 20h was calculated as a ratio ((w_0_-w_20_)/w_0_) expressed as a percentage.

20h after infection the intestine (duodenum to rectum) was removed and its weight was measured(w_i_). The intestine to carcass was calculated as a ratio (w_i_/(w_20_-w_i_)). The intestines were homogenized in 1ml PBS using a stainless steel bead (cat no 69989, QIAGEN, Netherlands) and a TissueLyser II (QIAGEN, Netherlands) Time: 8 min, Frequency 30/s. A serial dilution of the homogenized material was plated out onto blood agar (Substrat, Sahlgrenska Universitetssjukhuset, Sweden) and Thiosulfate-citrate-bile salts-sucrose agar plates (86348, Merck, Germany). The number of *Vibrio cholerae* CFU per intestine was calculated.

### Ethical statement

All animals were housed under specific-pathogen-free conditions and all treatments and procedures were performed in accordance with the Swedish Animal Welfare Act (1988:534) and the Animal Welfare Ordinance (1988:539). Approval for the study was given by the Ethical Committee for Laboratory Animals in Gothenburg, Sweden (Ethical number 81/2016).

In order to minimize suffering, animals that during the course of the experiments showed clinical signs of severe disease i.e. change of skin coloration from pink to blue or grey, labored breathing and persistent recumbency, were immediately euthanized.

### Statistical analysis

Statistical analysis and differential expression analysis of RNAseq data were performed using the DESeq2 R package(34).

For data from CTB production experiment under AKI conditions, 10 cultures of each strain were cultivated and each sample was analyzed twice by GM1 ELISA. The mean value for each culture were used for the Two-tailed Unpaired t-test giving the following results: p=0.0036, t=3.345, df=18.

For data from the infant mouse experiments significance levels have been calculating using One-way ANOVA with Tukey’s multiple comparisons test regarding p adjusted values less than 0.05 as significant. Total number of mice per group from 7 independent experiments: MS1571 (23), MS1712 (23), MS1712 10x dose (6), phil6973 (16), PBS (23), and A493 (41), MS1843 (32), MS1972(10), MS1843 x10 dose (12), PBS (23). Brown-Forsythe test: F (DFn, DFd) values for: MS1571 vs MS1712 Weight loss 0,7199 (5, 89), Intestine/Carcass 6,438 (5, 90), Number of VC 1,58 (4, 66). A493 vs MS1843 Weight loss 4,424 (3, 75), Intestine/Carcass F 7,485 (3, 112), Number of VC F 1,199 (2, 82).

Data were analyzed with Prism 7.03 (GraphPad Software inc.)

## Author contributions

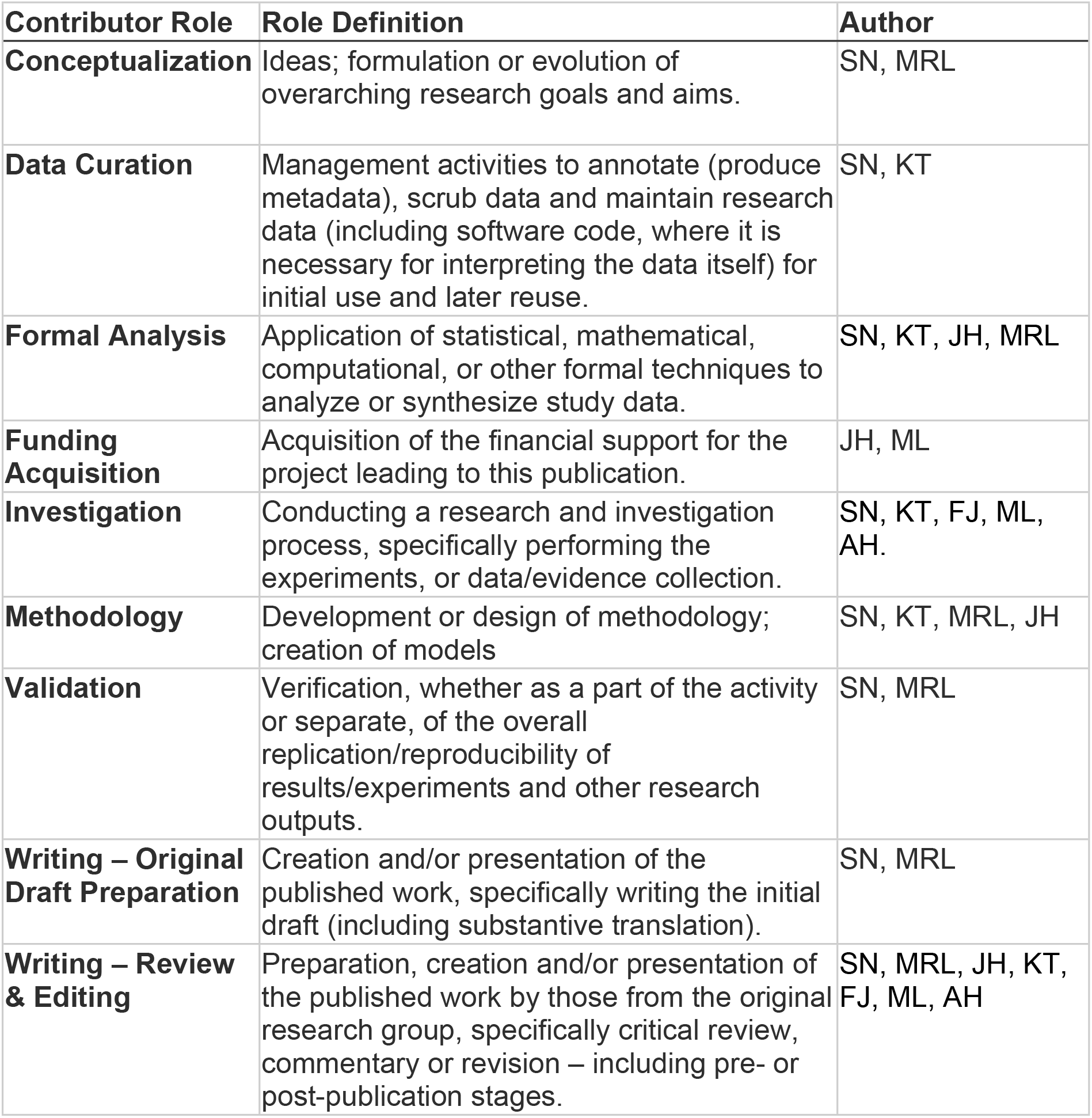

## Data availability

All data generated or analyzed during this study are either included in this published article (and its supplementary information files) or are available from the corresponding author on reasonable request.

## Supporting information captions

**Figure S1.** Direct comparison of growth of Ogawa and Inaba isogenic strains in LB growth media at 37°C

**Figure S2.** RNAseq Data, Pyrimidine metabolism

**Figure S3.** RNAseq Data, Purine metabolism

**Figure S4.** Infant Mouse Data.

**Figure S5.** Plasmid map for pMT-suicide1-sacB + wbeT + loxP + KmR

**Table S1** (Excel). RNAseq results: Genes significantly differentially expressed comparing strains MS1571 and MS1712 in mid-log phase in high salt medium at 24°C.

**Table S2** (Excel). RT-PCR primers

